# Mechanistic understanding enables the rational design of salicylanilide combination therapies for Gram-negative infections

**DOI:** 10.1101/2020.04.23.058875

**Authors:** Janine N. Copp, Daniel Pletzer, Alistair S. Brown, Joris Van der Heijden, Charlotte M. Miton, Rebecca J. Edgar, Michelle H. Rich, Rory F Little, Elsie M. Williams, Robert E.W. Hancock, Nobuhiko Tokuriki, David F. Ackerley

**Author notes:** Daniel Pletzer, Department of Microbiology and Immunology, University of Otago, New Zealand; Joris Van der Heijden, Ottawa Virus Manufacturing Facility, Ottawa Hospital Research Institute, Ottawa, Canada; Michelle Rich, School of Biomolecular and Biomedical Sciences, O’Brien Centre for Science, University College Dublin, Ireland.

## Abstract

One avenue to combat multidrug-resistant Gram-negative bacteria is the co-administration of multiple drugs (combination therapy), which can be particularly promising if drugs synergize. The identification of synergistic drug combinations, however, is challenging. Detailed understanding of antibiotic mechanisms can address this issue by facilitating the rational design of improved combination therapies. Here, using diverse biochemical and genetic assays, we reveal the molecular mechanisms of niclosamide, a clinically-approved salicylanilide compound, and demonstrate its potential for Gram-negative combination therapies. We discovered that Gram-negative bacteria possess two innate resistance mechanisms that reduce their niclosamide susceptibility: a primary mechanism mediated by multidrug efflux pumps and a secondary mechanism of nitroreduction. When efflux was compromised, niclosamide became a potent antibiotic, dissipating the proton motive force (PMF), increasing oxidative stress and reducing ATP production to cause cell death. These insights guided the identification of diverse compounds that synergized with salicylanilides when co-administered (efflux inhibitors, membrane permeabilizers, and antibiotics that are expelled by PMF-dependent efflux), thus suggesting that salicylanilide compounds may have broad utility in combination therapies. We validate these findings *in vivo* using a murine abscess model, where we show that niclosamide synergizes with the membrane permeabilizing antibiotic colistin against high-density infections of multidrug-resistant Gram-negative clinical isolates. We further demonstrate that enhanced nitroreductase activity is a potential route to adaptive niclosamide resistance but show that this causes collateral susceptibility to clinical nitro-prodrug antibiotics. Thus, we highlight how mechanistic understanding of mode of action, innate/adaptive resistance, and synergy can rationally guide the discovery, development and stewardship of novel combination therapies.

**Importance:** There is a critical need for more effective treatments to combat multidrug-resistant Gram-negative infections. Combination therapies are a promising strategy, especially when these enable existing clinical drugs to be repurposed as antibiotics. We reveal the mechanisms of action and basis of innate Gram-negative resistance for the anthelmintic drug niclosamide, and subsequently exploit this information to demonstrate that niclosamide and analogs kill Gram-negative bacteria when combined with antibiotics that inhibit drug efflux or permeabilize membranes. We confirm the synergistic potential of niclosamide *in vitro* against a diverse range of recalcitrant Gram-negative clinical isolates, and *in vivo* in a mouse abscess model. We also demonstrate that nitroreductases can confer resistance to niclosamide, but show that evolution of these enzymes for enhanced niclosamide resistance confers a collateral sensitivity to other clinical antibiotics. Our results highlight how detailed mechanistic understanding can accelerate the evaluation and implementation of new combination therapies.

## Introduction

New therapeutic strategies are urgently required to combat multidrug-resistant (MDR) Gram-negative bacteria (1). The co-administration of two or more drugs (combination therapy) (2) is a promising approach, especially if the drugs exhibit synergy, *i.e.*, enhanced efficacy over the predicted additive effects (3, 4). Synergistic combination therapies can kill microbes that are resistant to one of the drugs in the pair, may slow the evolution of resistance (5–7), and can facilitate the use of lower doses of each drug, thus reducing side effects and adverse reactions (8). The identification of synergistic drug combinations, however, is challenging due to the infrequency of synergistic relationships and the substantial scale of combinatorial drug screening (*e.g*., for a collection of 1,000 compounds, 499,500 pairwise combinations are possible, even before considering optimal relative concentrations) (3, 4). Mechanistic understanding of synergy may reveal novel antibiotic targets and guide the rational design of superior drug combinations *e.g.*, the co-administration of β-lactam compounds and β-lactamase inhibitors (9). However, for the majority of combination therapies the underlying basis of synergism is unclear. Indeed, despite clinical use for almost 50 years, the synergism of trimethoprim and sulfamethoxazole was only explored in detail in 2018 (10). Combination therapy may also enable compounds that have been clinically approved for other conditions, *e.g.*, antidepressants, antipsychotics, and antidiarrhetics (11–13), to be repurposed as antibiotics; such compounds typically have detailed data regarding their toxicity, formulation and pharmacology that can expedite their clinical progression (14). However, the screening of clinical compounds for repurposing potential is laborious and often necessitates high throughput robotic systems (13, 15). Detailed knowledge of the antibiotic mechanisms of action of promising clinical compounds would accelerate drug repurposing approaches and enable the circumvention of resistance mechanisms that may mask activity in initial screens. Thus, comprehensive understanding of both the mode of action and innate resistance mechanisms is important to inform the rational design of superior combination therapies that harness repurposed clinical compounds.

Niclosamide (**Fig. 1a**) is a clinically-approved drug that has been used to treat helminth parasites in humans and animals for more than 50 years (16). Recently, several studies have suggested the potential of repurposing niclosamide for other medical applications, *e.g*., niclosamide appears to modulate metabolic disorders and neurological conditions, and has antiproliferative effects against various cancers (17). The diverse pharmacological activities of niclosamide are likely the result of oxidative phosphorylation uncoupling and the modulation of signaling pathways (18, 19). Niclosamide exhibits antiviral activity against SARS-COV (20, 21), and is an effective antibiotic against Gram-positive and acid-fast pathogens (*e.g. Staphylococcus aureus and Mycobacterium tuberculosis*), as well as *Helicobacter pylori* (22–24). As an anti-infective, the low absorption and poor oral bioavailability of niclosamide may hamper its use (25), however optimized derivatives, nano-based formulations and/or local administration may rescue its therapeutic potential (26–28). In isolation, niclosamide exhibits no activity against most Gram-negative pathogens (1, 23). Nevertheless, it was recently reported that *in vitro* co-administration of niclosamide and colistin can overcome colistin-resistance in Gram-negative bacteria (29–31). While these findings suggest that niclosamide may hold cryptic antibiotic potential, the molecular basis that underlies its antibiotic mode of action, synergy, and the lack of efficacy against Gram-negative bacteria has been hitherto unknown.

**Fig. 1.**
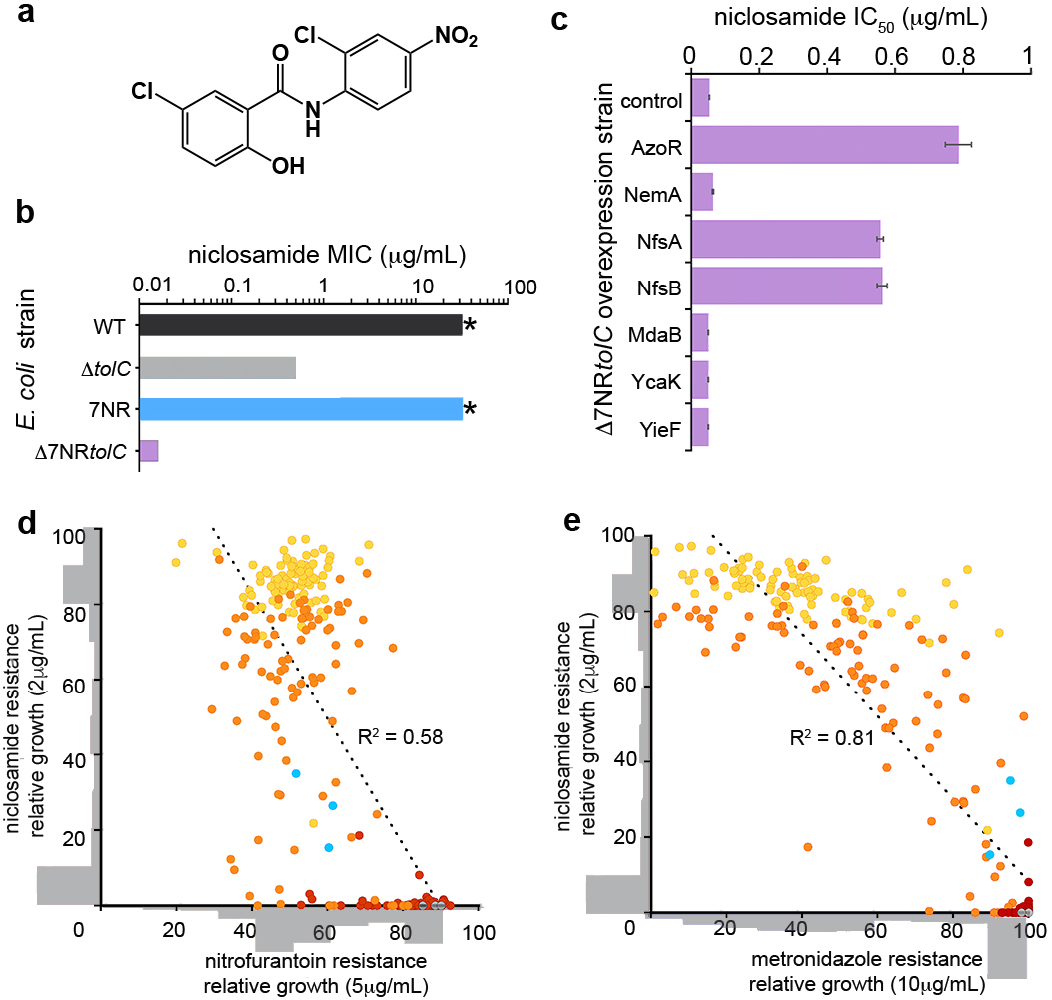
Niclosamide resistance mechanisms. **a**, Structure of niclosamide. **b**, MIC of *E. coli* strains: wild type (WT), Δ*tolC*, Δ7NR, and Δ7NR*tolC*; * indicates >32 μg.mL^−1^, which is the solubility limit of niclosamide in growth media. **c**, IC_50_ analysis of Δ7NR*tolC* strains individually overexpressing candidate *E. coli* nitroreductases or a vector only control following niclosamide administration. Error bars indicate SEM. **d-e**, Covariance plots displaying the interrelated profiles of niclosamide, metronidazole or nitrofurantoin resistance. 90 colonies of NfsA variants were picked from agar plates without niclosamide (red), 0.2 μg.mL^−1^ niclosamide (orange), or 2 μg.mL^−^ 1 niclosamide (yellow). Variants were grown overnight and then screened for niclosamide resistance (growth at 2 μg.ml^−1^) and (**d**) nitrofurantoin or (**e**) metronidazole resistance (growth at 5 and 10 μg.ml^−1^, respectively). Variant distribution is shown as grey histograms that are overlaid on the x and y axes. R^2^ values (linear regression analysis) are displayed; p <0.01. *E. coli* NfsA and vector-only controls are displayed in cyan and grey, respectively. All panels are constructed from pooled data from at least three independent biological replicates.

In this work, we uncover the innate resistance mechanisms and antibacterial mode of action of niclosamide and related salicylanilide analogs, thus revealing their therapeutic potential as potent antibiotics when utilized in rationally designed combination therapies. We reveal a potential route to adaptive niclosamide resistance, but demonstrate that this leads to collateral susceptibility; thus, the emergence of resistance via this route may be prevented or slowed in the clinic. In addition, we demonstrate the *in vitro* efficacy of niclosamide combination therapy against MDR Gram-negative clinical isolates, and confirm synergy *in vivo* using in a murine abscess model using high density infections that mimics a clinical situation where antibiotics typically fail (32).

## Results

### TolC-mediated efflux and nitroreductases confer innate niclosamide resistance in E. coli

To investigate the mechanisms by which *Escherichia coli* mitigates the antibiotic potential of niclosamide, we first examined multidrug efflux pumps, a dominant *E. coli* resistance mechanism to expel toxic compounds (33). We tested a variety of *E. coli* strains that lacked individual components of the three major tripartite efflux systems to ascertain whether efflux contributed to niclosamide resistance. In total, nine individual gene deletions were investigated for their effect on niclosamide minimal inhibitory concentration (MIC) (Table S1). Notably, deletion of the gene encoding the outer membrane channel TolC (Δ*tolC*) reduced MIC by >64-fold (MIC = 0.5 μg.mL^−1^), whereas no other deletions had any effect (MIC >32 μg.mL^−1^; Table S1). This result suggests that TolC-mediated efflux is one of the predominant mechanisms of niclosamide resistance. Interestingly, deletion of genes encoding other components of the principal RND-type TolC tripartite complex (AcrA or AcrB) had no effect on MIC. This was likely due to TolC interacting with alternative efflux components such as AcrE or AcrF, resulting in alternative niclosamide-capable pump assemblies (33).

Next, we explored the role of azo- and nitro-reductase flavoenzymes in niclosamide susceptibility, due to their importance in diverse metabolic pathways including antibiotic metabolism (34, 35). Although previous antibiotic metabolism studies have primarily focused on the bioreductive activation of nitro-prodrugs, we considered there was potential for nitroreduction to here be a detoxifying mechanism, as there is evidence that the nitro-moiety of niclosamide is an important structural feature for uncoupling activity (36). To test this, we generated an *E. coli* strain that lacked seven flavoenzyme genes with confirmed or putative nitro- or azo-reductase activity (Δ7NR) (37) (Table S1). Although niclosamide resistance in Δ7NR did not change compared to wild type *E. coli* (MIC >32 μg.mL^−1^), an otherwise isogenic strain that also lacked TolC (Δ7NR*tolC*) was 2000-fold more susceptible to niclosamide than wild type (MIC = 0.016 μg.mL^−1^) and 32-fold more susceptible than Δ*tolC* (**Fig. 1b**; Table S1). The relative contributions of each of the seven flavoenzymes were delineated by individually overexpressing the corresponding genes in Δ7NR*tolC* and investigating their effects on niclosamide IC_50_ (the concentration of niclosamide required to reduce the bacterial burden by 50%). We demonstrated that three enzymes, NfsA, NfsB, and AzoR, increased niclosamide IC_50_ by 10-to 15-fold (**Fig. 1c**). Although these three enzymes derive from two distinct structural folds, they are all proficient nitroreductases (37). Increasing nitroreductase activity could therefore be a potential adaptive strategy for bacteria to develop resistance against niclosamide. We hypothesized, however, that this might cause collateral sensitivity to nitroaromatic prodrug antibiotics such as nitrofurantoin or metronidazole. To test this hypothesis, we selected Δ7NR*tolC* cells expressing different NfsA variants (generated via multisite saturation mutagenesis to combinatorically randomize seven active site residues) for resistance to either 0.2 or 2μg.mL-1 niclosamide, then counter screened for sensitivity to nitrofurantoin or metronidazole. Consistent with our hypothesis, increasing niclosamide resistance via more proficient nitroreductase variants concomitantly decreased nitrofurantoin and metronidazole resistance (**Fig. 1d-e**).

### Niclosamide disrupts oxidative phosphorylation in E. coli

Next, the underlying mechanisms of niclosamide antibiotic activity were investigated. As previous studies have demonstrated that niclosamide uncouples oxidative phosphorylation in mitochondria and *H. pylori* (18, 24), multiple physiological attributes were explored that relate to this process in Gram-negative bacteria, namely proton motive force (PMF), oxygen consumption, ATP production, and redox homeostasis. The PMF has two parameters: the electric potential (ΔΨ) and the transmembrane proton (ΔpH) gradients. First, the effect of niclosamide on PMF-dissipation was investigated in EDTA-permeabilized *E. coli* using a fluorescent assay that employs the membrane potential sensitive dye diSC_3_(5) - a caged cation that distributes in the membrane according to ΔΨ and self-quenches. We observed that niclosamide specifically dissipated the ΔΨ (Fig. 2a) as revealed by de-quenching of diSC_3_5 fluorescence.

**Fig. 2.**
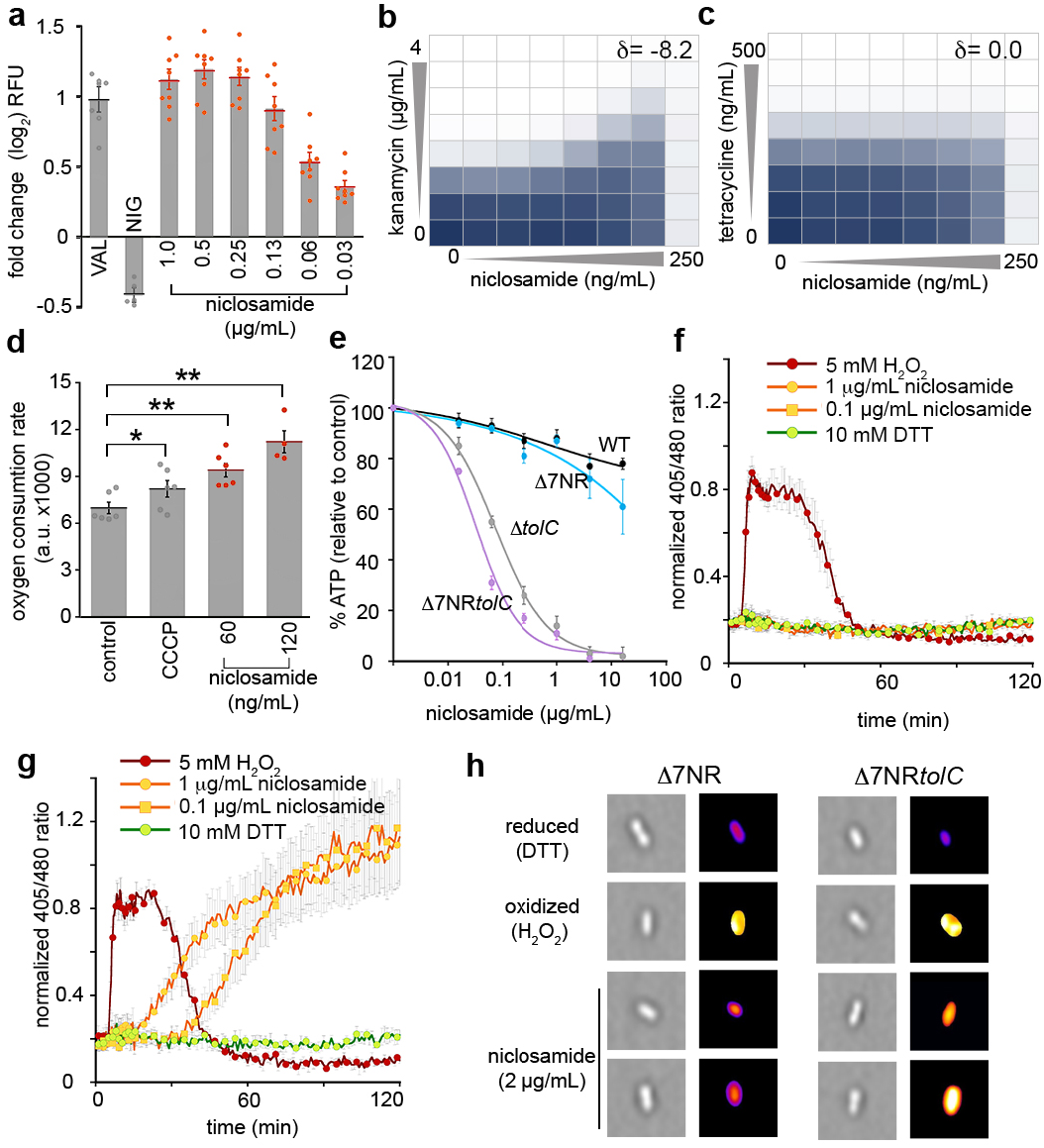
Antibiotic mechanisms of niclosamide. **a**. Fold change in diSC_3_(5) fluorescence. *E. coli* was grown in MHB with 10 mM EDTA to an OD_600_ of 1. Cells were incubated with diSC_3_(5) for 20 min prior to administration of 0.5 μg/mL valinomycin (VAL; a ΔΨ-dissipating ionophore), 0.5 μg/mL nigericin (NIG; a ΔpH-dissipating ionophore) or 0.03 to 1 μg/mL niclosamide. 100 mM KCl was added to cells prior to valinomycin treatment. **b**-**c**, The combined inhibitory effects of 0 - 250 ng.mL^−1^ niclosamide and either (**b**) 0 - 4 μg.mL^−1^ kanamycin, or (**c**) 0 - 500 ng.mL^−1^ tetracycline were tested against Δ7NR*tolC* in a checkerboard format. Bacterial growth is shown as a heat plot. **d**, Oxygen consumption was measured using the MitoXpress oxygen probe in Δ7NR*tolC* cells (mid-log; OD_600_ = 0.15) overlaid with mineral oil for 20 min. **e**, Relative cellular ATP levels were estimated by luciferase activity and compared to unchallenged (DMSO-only) control. **f-g**, Intracellular oxidation levels were measured in (**f**) WT *E. coli* and (**g**) Δ*tolC* strains constitutively expressing redox-sensitive GFP (roGFP) following administration of 5 mM H_2_O_2_ (oxidized control), 10 mM DTT (reduced control), or niclosamide. **h**, Representative high-throughput fluorescence microscopy images of Δ7NR and Δ7NR*tolC* cells 120 min after administration of DTT, H_2_O_2_, or niclosamide. Images on the right are pseudo-colored ratio images after analysis with ImageJ. Panels a-g were constructed from pooled data from at least three independent biological replicates. Labels indicate significant responses over the control (* = P<0.05; ** = P<0.01). Statistical analyses were performed using One-way ANOVA, Kruskal-Wallis test. Error bars indicate SEM.

To confirm this result, checkerboard assays (dose response growth assays using serial dilutions of two drugs in combination) were performed in Δ7NR*tolC* using niclosamide and antibiotics that rely upon either ΔΨ or ΔpH for cell uptake (kanamycin and tetracycline, respectively). The fractional inhibitory concentration index is frequently used to characterize drug interactions, but has limitations when analyzing compounds for which an individual MIC cannot be obtained (here, niclosamide). Thus, drug interactions were analyzed via Zero Interaction Potency (ZIP) scores (δ) that quantify the change in dose– response curves between individual drugs and combinations thereof, from the expectation of no interaction; δ scores >0 indicate synergism, 0 indicates no interaction, and <0 antagonism (38). Kanamycin efficacy was reduced in the presence of niclosamide, *i.e.*, niclosamide was antagonistic when co-administered with kanamycin, corresponding to a δ score of −8.2 (**Fig. 2b**), which is consistent with ΔΨ dissipation undermining ΔΨ-dependent kanamycin uptake. In contrast, tetracycline efficacy was not affected by niclosamide co-administration (δ = 0.0; Fig. 2c), as tetracycline relies upon ΔpH for uptake (**Fig. 2b-c**). PMF disruption can reduce ATP production and increase both oxygen consumption and oxidative stress (39). We therefore confirmed that niclosamide administration significantly increased oxygen consumption in Δ7NR*tolC* (by 1.4- and 1.6-fold after administration of 60 ng.mL^−1^ and 120 ng.mL^−1^ niclosamide, respectively; **Fig. 2d**). Niclosamide administration caused a reduction of cellular ATP concentration to 3% and 1% of DMSO-control concentrations when 4 μg.mL^−1^ niclosamide was administered to Δ*tolC* and Δ7NR*tolC*, respectively (**Fig. 2e**). Employing strains constitutively expressing redox-sensitive GFP (40), we determined that niclosamide also disrupted redox homeostasis in Δ*tolC* and Δ7NR*tolC* strains, causing an increase in oxidative stress (**Fig. 2f-g**; Fig. S1). Next, using high-throughput fluorescence microscopy, increased intracellular oxidative stress was visualized in Δ7NR*tolC* cells following niclosamide administration. The distribution of redox stress per cell was plotted as histograms (Fig. S1) and a random selection of pseudo-colored ratio images are presented in **Fig. 2h**. In strains that retained TolC function, niclosamide did not have a significant effect on cellular ATP levels, oxygen consumption, or redox homeostasis. Taken together, these data suggest that, when efflux is compromised, niclosamide dissipates the ΔΨ to collapse the PMF and uncouple oxidative phosphorylation in *E. coli*.

### Niclosamide synergizes with efflux pump inhibitors and membrane permeabilizers for enhanced efficacy against E. coli

After establishing the mode of action of niclosamide and the basis of Gram-negative innate resistance, we next sought to identify compounds that sensitize Gram-negative bacteria to niclosamide when administered in combination. Predicting that efflux pump inhibitors such as phenylalanine-arginine β-naphthylamide (PAβN) (41) would increase niclosamide sensitivity, checkerboard assays were employed to screen niclosamide and PAβN against *E. coli*. Considerable synergy was observed (δ = 47.9; **Fig. 3a**). It was next reasoned that increased niclosamide influx via outer membrane permeabilization might mitigate TolC-mediated efflux. Therefore, membrane permeabilizing polymyxin antibiotics were investigated for synergy and, consistent with recent reports (29–31), synergy was observed when colistin or polymyxin B were co-administered with niclosamide (δ = 28.1 and 25.6, respectively) (**Fig. 3b-c**). We hypothesized that synergism was due to the cascading effect of the mode of action of niclosamide, in that polymyxins increased the influx of niclosamide and thereby facilitated PMF dissipation, which in turn compromised the efficiency of PMF-dependent niclosamide efflux (as efflux was dependent upon PMF (42)). Ultimately, this would result in higher intracellular concentrations of niclosamide and thus enhanced antibiotic effects (Fig. S2). Indeed, polymyxin synergy was less evident in Δ*tolC* (δ = 7.6 and 9.7 for colistin and polymyxin B respectively; **Fig. 3d-e**) and niclosamide administration inhibited efflux in EDTA-permeabilized *E. coli* (observed via increasing intracellular accumulation of the fluorescent nucleic acid probe Hoechst 33342; Fig. S1).

**Fig. 3.**
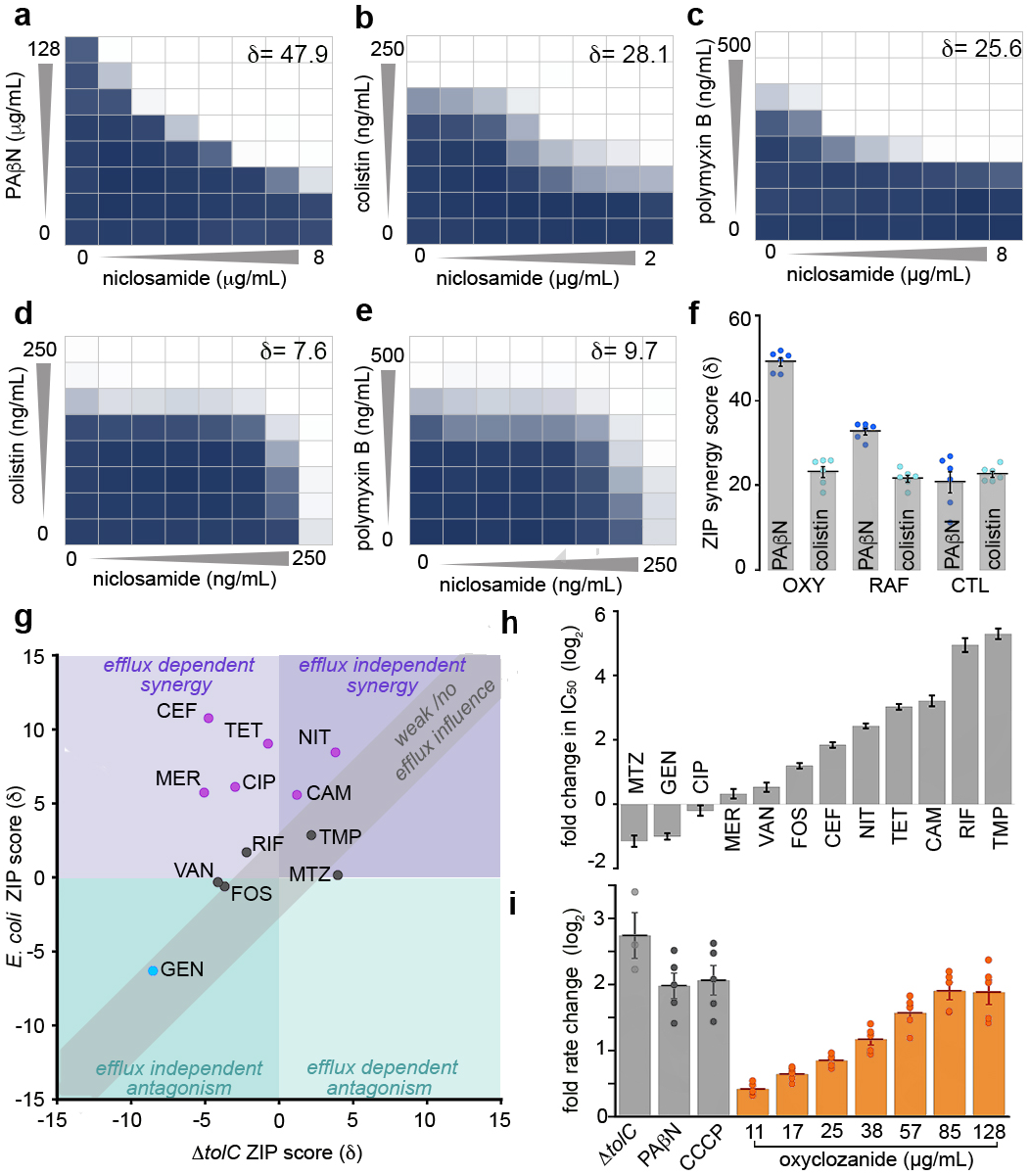
Analyses of salicylanilide synergy interactions. **a-c**, The combined inhibitory effects of 0 - 8 μg.mL^−1^ niclosamide and either (**a**) 0 - 128 μg.mL^−1^ PAβN, or (**b**) 0 - 250 ng.mL^−1^ colistin, or (**c**) 0 – 500 ng.mL^−1^ polymyxin B were tested against *E. coli* using checkerboard analysis. ZIP synergy scores (δ) are presented. Bacterial growth is depicted as a heat plot. **d-e**, The combined inhibitory effects of 0 to 250 ng.mL^−1^ niclosamide and either (**d**) 0 to 250 ng.mL^−1^ colistin or (**e**) 0 – 500 ng.mL^−1^ polymyxin B were tested against *E. coli* Δ*tolC* in checkerboard analyses. Bacterial growth is depicted as a heat plot. **f**, A bar graph of ZIP scores (δ) depicting the synergism of oxyclozanide (OXY), rafoxanide (RAF), or closantel (CTL) in combination with PAβN or colistin against *E. coli*. Error bars indicate SEM. **g-h**, Analysis of oxyclozanide synergy with: nitrofurantoin (NIT), metronidazole (MTZ), cefotaxime (CEF), rifampicin (RIF), tetracycline (TET), gentamicin (GEN), ciprofloxacin (CIP), chloramphenicol (CAM), trimethoprim (TMP), fosfomycin (FOS), meropenem, (MER) or vancomycin (VAN). (**g**) A covariance plot of antibiotic ZIP scores from checkerboard assays conducted in minimal media with oxyclozanide against *E. coli* and Δ*tolC*. (**h**) A bar chart displaying the fold change of IC_50_ values in *E. coli* compared to Δ*tolC*; uncertainty is indicated by error bars. **i**, Fold change in rate of H33342 fluorescence (compared to a DMSO control) in Δ*tolC* cells or WT *E. coli* following administration of 28 μg.mL^−1^ PAβN, 5 μg.mL^−1^ CCCP, or 11.2 to 128 μg.mL^−1^ oxyclozanide. Error bars indicate SEM. All panels were constructed from pooled data from at least three independent biological replicates.

### Oxyclozanide potentiates diverse antibiotics, likely via inhibition of PMF-dependent efflux

Next, it was examined whether the synergistic relationships observed above were maintained for other halogenated salicylanilides, namely oxyclozanide, closantel, and rafoxanide. It was confirmed that all these niclosamide analogs synergized with both PAβN and colistin (δ = 21.5 to 49.8; **Fig. 3f**; Fig. S3). Of note, the relatively high solubility of oxyclozanide in growth media (~512 μg.mL^−1^), compared to that of other salicylanilides (~64 μg.mL^−1^), enabled the observation of an oxyclozanide MIC (256 μg.mL^−1^), *i.e.*, sufficiently high concentrations of oxyclozanide could overcome TolC-mediated efflux. Since moderate synergistic relationships can only be detected at concentrations nearing the MIC of both drugs and are emphasized in bacterial cultures under nutrient limitation (43), oxyclozanide checkerboard assays in minimal media were used to identify additional antibiotics that interact with salicylanilides against *E. coli*. Twelve antibiotics with diverse cellular targets (Table S1) were examined. Interestingly, 6 out of 12 antibiotics synergized with oxyclozanide (chloramphenicol, tetracycline, cefotaxime, meropenem, ciprofloxacin, and nitrofurantoin; δ = 5.7 to 10.8), five antibiotics displayed no or weak interactions (δ = −0.6 to 2.9) and, consistent with the dependence of aminoglycosides on ΔΨ for uptake, oxyclozanide antagonized gentamicin activity (δ = −6.3) (**Fig. 3g**; Table S2; **Fig. 4**). These combinatorial effects were neutralized or reversed when examined in the Δ*tolC* strain (**Fig. 3g**; Table S2; **Fig. 4**) and oxyclozanide synergy was typically stronger for antibiotics that are TolC-substrates (indicated by fold IC_50_ change in Δ*tolC* compared to *E. coli*; **Fig. 3h**). These results suggest that oxyclozanide synergy might be at least partially underpinned by the inhibition of efflux via PMF dissipation (Fig. S2). To substantiate this hypothesis, the effect of oxyclozanide on cellular efflux was examined using Hoechst 33342. Indeed, administration of oxyclozanide inhibited efflux (**Fig. 3i**). Taken together, these results demonstrate that chemical disruption of TolC-mediated efflux or membrane integrity sensitizes *E. coli* to salicylanilides. Further studies are required to explain detailed drug interactions, however, our results suggest that PMF-dissipating compounds such as salicylanilides may potentiate the activity of diverse antibiotics through the disruption of efflux.

**Fig. 4.**
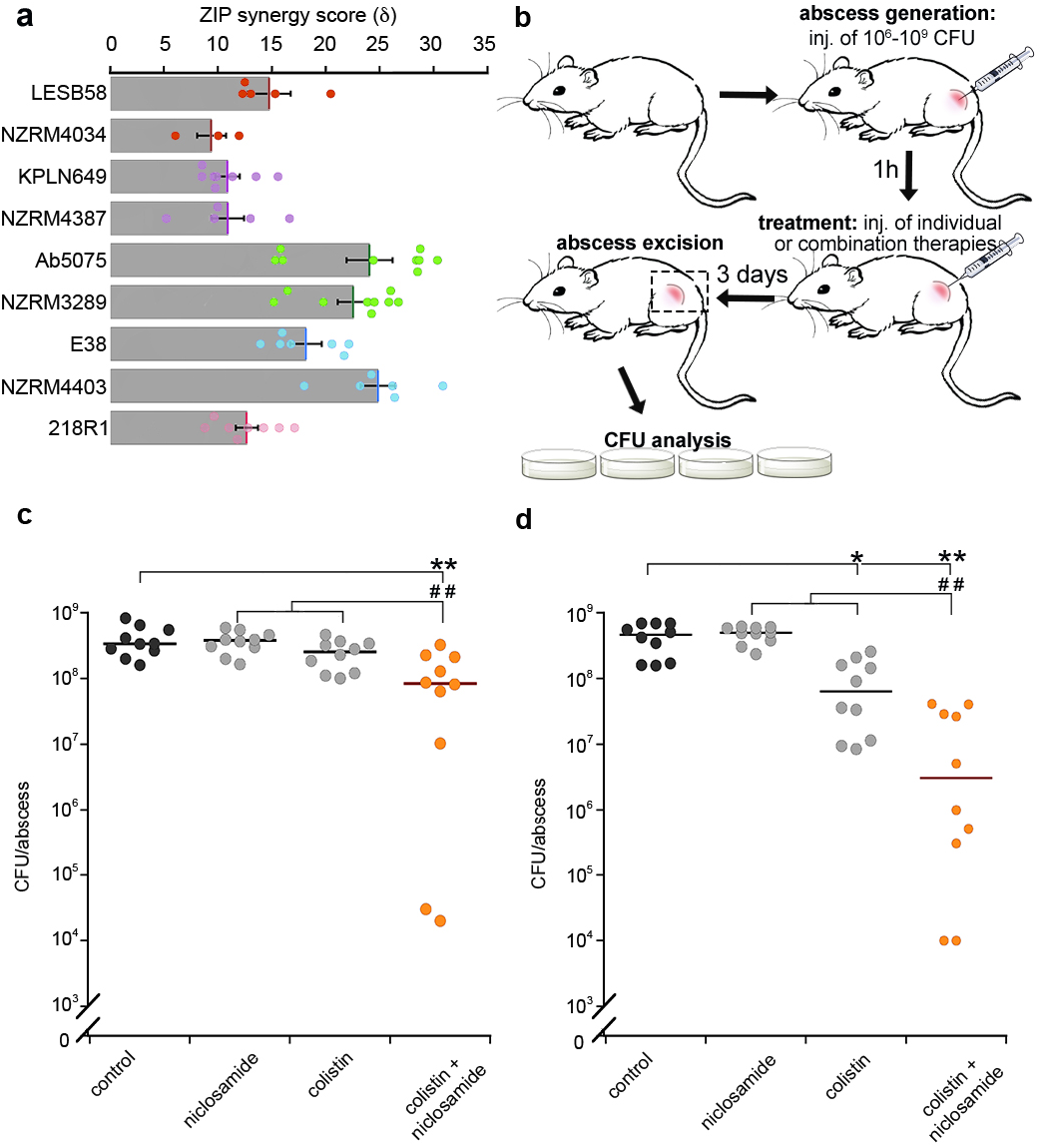
Niclosamide/colistin combination therapy was effective against recalcitrant MDR Gram-negative strains. **a**, Bar graph depicting *in vitro* ZIP scores (δ) of niclosamide and colistin co-administration against clinical MDR Gram-negative strains: *P. aeruginosa* LESB58, *P. aeruginosa* NZRM4034, *K. pneumoniae* KPLN649, *K. pneumoniae* NZRM4387, *A. baumannii* Ab5075, *A. baumannii* NZRM3289, *E. coli* E38, *E. coli* NZRM4403 and *E. cloacae* 218R1. The ZIP synergy score (δ) is presented as the average interaction from an 8×8 dose-response matrix. Data was averaged from at least three independent experiments and error bars indicate SEM. **b**, Diagram of abscess model procedure and analysis. **c-d**, Dot plots of (**c**) colistin-resistant *P. aeruginosa* LESB58 and (**d**) *K. pneumoniae* KPLN649 survival, represented as CFUs recovered per abscess after administration of 10 mg.kg^−1^ niclosamide ethanolamine salt and 0.15 mg.kg^−1^ (*P. aeruginosa*) or 2.5 mg.kg^−1^ (*K. pneumoniae*) colistin as individual or combined therapeutics. Labels indicate: significant responses over the PEG control (*, p <0.05; **, p <0.01); or synergistic responses, *i.e.*, significant differences of the combination therapy over the sum of the effects of each agent alone (^##^, p <0.01). Statistical analyses were performed using One-way ANOVA, Kruskal-Wallis test with Dunn’s correction (two sided).

### Niclosamide/colistin combination therapy is effective against diverse clinical Gram-negative isolates in vitro and in vivo

Finally, we investigated the potential of salicylanilide combination therapy against a range of Gram-negative clinical isolates. Due to the immediate repurposing potential of niclosamide as an FDA-approved clinical drug and increasing concerns around colistin-resistant pathogens, we prioritized these two compounds. Checkerboard assays were performed on nine MDR clinical isolates across diverse phyla: *Acinetobacter*, *Pseudomonas* and the *Enterobacteriaceae* (Table S3). Co-administration of niclosamide and colistin yielded synergistic efficacy in all strains including the colistin resistant clinical isolate, *P. aeruginosa* LESB58 (δ = 8.4 to 36.7), enabling up to 4-fold lower doses of colistin (**Fig. 4a;** Fig. S5), which is of particular importance due to the nephrotoxicity issues associated with this antibiotic (44).

The poor bioavailability and pharmacology of niclosamide may be mitigated via local administration, *e.g*., topical or inhalation therapies (28, 45). Here, we examined the *in vivo* antibacterial synergy of niclosamide via direct injection in a high-density murine cutaneous infection model that mimics clinical situations where antibiotic treatments are typically unsuccessful, *e.g.*, skin abscesses (32) (**Fig. 4b**). The synergistic efficacy of niclosamide and colistin was validated against *P. aeruginosa* LESB58 and *K. pneumoniae* KPLN649. Co-administration resulted in significant synergistic efficacy against both strains, reducing the *K. pneumoniae* and *P. aeruginosa* bacterial load by 32- and 12-fold, respectively, over the sum of the individual therapies, and by 239- and 19-fold, respectively, when compared to vehicle-only controls (**Fig. 4c-d**). This is the first report of *in vivo* efficacy for niclosamide and colistin against Gram-negative pathogens, and notably, this was achieved against recalcitrant high-density infections for which no effective clinical treatments currently exist (32). It is also important to note that this study was focused on detecting niclosamide/colistin synergy rather than identifying the best formulation or dose ratio for efficacy; more significant efficacy could likely be achieved by optimising the drug concentrations or dosing regimen.

## Discussion

By harnessing a diverse set of biochemical and genetic tools, this work explores the Gram-negative antibacterial potential of niclosamide and related salicylanilide analogs. We reveal the molecular action of salicylanilides against Gram-negative bacteria, detailing not only the underlying mechanisms of antibiotic activity, but also the basis for innate and adaptive resistance and the mechanisms that underpin the synergies between salicylanilides and a diversity of other antibiotics. These data enabled the development of a model that substantially advances our knowledge of the physiological effects of salicylanilides in Gram-negative bacteria (Fig. S2). Efflux is an established Gram-negative resistance mechanism and indeed, this is the predominant basis for niclosamide resistance. However, we demonstrate that salicylanilides also inhibit efflux and thus synergize with a wide range of antibiotics for which efflux is a common resistance mechanism. These data highlight the potential of the salicylanilide chemical scaffold, and PMF-dissipating compounds in general, for the discovery and design of novel antibiotic adjuvants to address efflux-mediated resistance. We consider PMF-dissipation, traditionally avoided in early drug development due to presumed toxicity, a promising and unexplored trait for the development of antimicrobials (46). Interestingly, many clinical compounds for diverse medical purposes have been reported to have mild PMF-dissipating activity and in addition, some have displayed antibiotic efficacy against Gram-positive or acid-fast pathogens such as *S. aureus* and *M. tuberculosis* (46, 47). Gram-negative pathogens, in contrast resist the action of such compounds via their robust cellular envelope and diverse efflux pumps (33). Co-administration of PMF-dissipating compounds with drugs that target efflux and cell permeability may therefore be a promising avenue to discover more effective combination therapies. Combining mechanistic insights with previously established data around safety and pharmacology for repurposed “non-antibiotic” clinical compounds may rapidly identify attractive candidates for accelerated clinical development.

Understanding the evolutionary basis of antibiotic resistance is important to inform the sustainable use of next generation antibiotics. Due to the failure of laboratory evolution experiments to generate niclosamide resistance in *S. aureus* or *H. pylori*, previous reports have suggested that a key advantage of niclosamide as a potential antimicrobial is the apparent lack of resistance mechanisms (24, 45). We show, however, that nitroreductases inactivate niclosamide to reduce antibiotic toxicity and enhanced nitroreductase activity can cause niclosamide resistance. While modulation of nitroreductase activity is a known Gram-negative resistance mechanism against nitro-antibiotic compounds, this is typically caused by null mutations, *i.e.*, a genetic change that results in a non-functional nitroreductase, to prevent the activation of prodrug antibiotics such as metronidazole (48). Our results suggest that resistance has potential to emerge in the clinic through enhanced nitroreductase activity. Significantly, we show that this may result in collateral sensitivity to nitroimidazole antibiotics, and thus propose a strategy to mitigate this evolutionary route, *i.e.* cyclic treatments of metronidazole. This demonstrates how mechanistic understanding can accelerate not only the discovery, but also potentially the sustainability of new Gram-negative combination therapies.

In summary, we reveal the detailed mechanisms that underlie the antibiotic mode of action, routes of resistance and synergistic relationships of salicylanilides. This guided the discovery of novel combination therapies and emphasizes how mechanistic understanding is critical when seeking to repurpose clinical compounds. Salicylanilides, and likely other PMF-dissipating compounds, may have broad utility as Gram-negative antibiotics in combination therapies.

## Acknowledgements

This work was supported by the Royal Society of New Zealand Marsden Fund (grant VUW1502 to D.F.A.), the Health Research Council of New Zealand (18-532 to D.F.A.), a Natural Sciences and Engineering Research Council of Canada Discovery Grant (RGPIN 2017-04909 to N.T.) and a Canadian Institutes for Health Research (CIHR) grant (FDN-154287 to R.E.W.H.). D.P. received a Cystic Fibrosis Postdoctoral Fellowship (Canada) and a Research Trainee Award from the Michael Smith Foundation for Health Research. R.E.W.H. holds a Canada Research Chair in Health and Genomics and a UBC Killam Professorship. N.T. is a CIHR new investigator and a Michael Smith Foundation of Health Research career investigator. J.N.C. thanks Jaiten Saini for technical support.

## Author contributions

J.N.C. and D.F.A. designed and directed the project, and co-wrote the manuscript. J.N.C performed the mechanistic studies and bacterial screening assays. N.T. assisted project design and co-wrote the manuscript. J.V.D.H. performed the AMNIS high throughput screening and assisted intracellular redox assays; M.H.R., A.S.B., R.F.L., C.M.M. and R.J.E. assisted bacterial screening; E.M.W. provided experimental support for molecular engineering and bacterial assays; D.P. carried out all abscess infection studies; D.P and R.E.W.H. directed the mouse abscess infection studies and assisted in manuscript editing.

## Methods

### Bacterial Strains

*E. coli* BW25113 strains bearing individual gene deletions were obtained from the Keio knockout collection (49). Δ7NR and Δ7NR*tolC* were generated via sequential knockout as previously described (50). New Zealand clinical isolates used in this study were *A. baumannii* NZRM3289, *P. aeruginosa* NZRM4034, *K. pneumoniae* NZRM4387 and *E. coli* NZRM4403 (obtained from the New Zealand Reference Culture Collection, Environmental Science and Research Ltd.), additional strains, *K. pneumoniae* KPLN649, *A. baumannii* Ab5075, *P. aeruginosa* LESB58, *E. cloacae* 218R1, *E. coli* E38 (Serotype O78:H^−^) (BEI resources, NR-17717) as previously described (51).

### In vitro growth analyses

Minimal inhibitory concentrations (MIC) were determined using two-fold dilutions and growth was measured after 16 - 48 h (52). The MIC was the concentration that inhibited growth >90% when compared to controls. DMSO was present at a final concentration of 2.5 % unless otherwise stated. For checkerboard analysis, an 8 × 12 matrix was created with two-fold serial dilutions of each compound. Bacterial colonies were isolated from a freshly streaked plate and resuspended in MHB media for OD_600_ normalization. After addition of bacteria to a final OD_600_ of 0.001, checkerboard plates were incubated at 30 °C with shaking for 16 h (or 36 h at 37 °C for *P. aeruginosa* strains), at which time the OD_600_ was measured. Checkerboard assays in minimal media were typically performed at 37 °C (oxyclozanide checkerboard assays with nitrofurantoin, rifampicin, tetracycline and chloramphenicol were performed at 30 °C) with shaking for 16 h, from a starting OD_600_ of 0.04. For the analysis of nitroreductase overexpression strains, individual colonies were transferred via nitrocellulose membrane to an agar plate containing 1mM IPTG and incubated for 3.5 h. IPTG-induced cells were then removed from the membrane and resuspended in MHB media for checkerboard analysis as described above. Relative IC_50_ values (the concentration of the compound required to reduce the bacterial burden by 50% compared to unchallenged controls) were calculated from the dose-response curves using the four-parameter equation y=m1+(m2-m1)/(1+(x/m3)^m4) determined by Kaleidagraph software (Synergy Software, Reading, PA) where m1 = lower asymptote, m2 = lower asymptote, m3 = relative IC_50_, and m4 = slope.

### Generation and screening of mutagenized NfsA variants

A plasmid-based multisite saturation mutagenesis library of *E. coli* NfsA (UniProtKB ID: P17117) was generated via combinatorial randomization of the codons encoding seven key active site residues: S41, F42, F83, K222, S224, R225 and F227 (Fig. S6). All codons were randomized to NDT (a degeneracy that specifies a balanced range of 12 different amino acids including the native residue), with the exception of position 222, which was randomized to NNK (specifying all 20 possible amino acids). The resulting library of nearly 96 million codon variants was expressed in Δ7NR*tolC*. To analyze the activity of NfsA variants, library subsets were selected on agar plates using 0, 0.2 and 2 μg.mL^−1^ of niclosamide. Ninety colonies from each subset were subsequently screened via growth assays with niclosamide, metronidazole, and nitrofurantoin at 2, 10, and 5 μg.mL^−1^ respectively; growth was measured via OD_600_ following 4 h incubation as previously described (53).

### Synergy Calculations

For each checkerboard analysis, an 8 × 8 matrix of averaged checkerboard results from at least three (typically >5) independent experiments was used to calculate ZIP scores using SynergyFinder (Bioconductor.org) (38, 54).

### DiSC_3_(5) assay

Subcultures of *E. coli* BW25113 were grown to late exponential phase (OD_600_ ~1) in MHB with 10 mM EDTA (to facilitate diSC_3_(5) cell entry). Cells were harvested by centrifugation, washed twice in buffer (5 mM HEPES, pH 7.2, 20 mM glucose, 5% DMSO), and then resuspended in buffer to a final OD_600_ = 0.085 with 1 μM DiSC3(5). For valinomycin, 100 mM KCl was added to the cell suspension containing diSC_3_(5). After a 20 min incubation at 37 °C, 190 μL of diSC_3_(5) loaded cells were added to two-fold dilutions of niclosamide, valinomycin, or nigericin in 96-well black clear-bottom plates (Corning, NY). Fluorescence (Ex: 620 nm, Em: 685 nm) was immediately read using a Synergy H1 Hybrid plate reader.

### Measurement of intracellular ATP levels

*E. coli* BW25113 was grown in MHB to early-log phase (OD_600_ = 0.2) and then grown in the presence of niclosamide or CCCP for 60 min in clear flat-bottom 96-well plates. The OD_600_ was determined immediately before ATP levels were measured using BacTiter-Glo™ (Promega, Madison WI), according to manufacturer instructions. Relative ATP levels were calculated by dividing relative light units (RLU) by the OD_600_ (RLU/OD).

### Measurement of Oxygen consumption

*E. coli* strains were grown in MHB to early-log phase (OD_600_ = 0.4) before dilution to OD_600_ = 0.1 prior to the assay. 50 μL of diluted culture was added to individual wells of a 96-well black clear-bottom plate (Corning, NY) containing 5 μL of a DMSO control, CCCP, or niclosamide at the desired concentration, and 5 μL of the MitoXpress oxygen probe. Cells were immediately covered with a layer of high-sensitivity mineral oil (50 μL) to restrict oxygen diffusion. Fluorescence (Ex: 380 nm, Em: 650 nm) was recorded using a Synergy H1 Hybrid plate reader.

### Measurement of bacterial efflux

Subcultures of *E. coli* BW25113 were grown to early exponential phase (OD_600_ ~0.4) in MHB supplemented with 5 mM EDTA, harvested by centrifugation, and resuspended in PBS to a final OD_600_ = 0.1. To initiate accumulation assays, Hoescht 33342 was added (1μM), cells were mixed by inversion and 150 μl aliquots were added in a black, clear-bottom 96-well plate containing 50 μL of oxyclozanide, niclosamide, PAβN or CCCP at 4× the desired concentration(s) in PBS with 20% DMSO. Fluorescence (Ex: 355 nm, Em:460 nm) was measured for 10 min using a Synergy H1 Hybrid plate reader (55).

### Measurement of intrabacterial redox potential

roGFP contains an intramolecular disulfide bond that induces a shift in fluorescence emission between 405 nm and 480 nm, thus intracellular oxidative stress can be ratiometrically monitored. *In vitro* analysis of the intrabacterial redox potential was performed as previously published (40) Assays were performed at 30 °C in a Synergy H1 Hybrid plate reader with excitation measured at 405 and 480 nm, and emission at 510 nm. Log phase bacterial cultures were resuspended in 0.9% sodium chloride at OD_600_ of 1.0, and 180 μL per well were loaded in a black, clear-bottom 96-well plate. The signals for fully oxidized or fully reduced bacteria were obtained by adding 5 mM H_2_O_2_ or 10 mM DTT to the bacteria culture at the start of the experiment. Niclosamide was added at 1 and 0.1 μg.mL^−1^. All values were normalized to the values obtained for maximally oxidized and for fully reduced bacterial cultures.

### AMNIS ImageStream and IDEAS/ImageJ Analysis

Samples were analysed by the AMNIS ImageStream as previously described (40). The laser intensities for 405, 488, 658, and 785 nm were 100, 120, 20, and 3.8, respectively. The data files were further analysed with the IDEAS software, version 6.0.129.0, which is supplied by AMNIS. Bacterial cells were selected based on fluorescence at 660 nm. Every cell image was then selected by the program based on fluorescent intensity at 660 nm. A mask was then created for analysis of the 405/480 nm ratio. The resulting 405/480 signals were plotted in a histogram. Reduced and oxidized controls were obtained within each experiment, niclosamide was administered at 2 μg.mL^−1^. All values were normalized to oxidized and reduced ratio values. Pseudo-colored ratio images were generated by ImageJ as described previously (40).

### Murine abscess infection studies

Animal experiments were performed in accordance with The Canadian Council on Animal Care (CCAC) guidelines and were approved by the University of British Columbia Animal Care Committee (certificate number A14-0363). Mice used in this study were female outbred CD-1. All animals were purchased from Charles River Laboratories (Wilmington, MA), were 7 weeks of age, and weighed 25 ± 3 g at the time of the experiments. 1 to 3% isoflurane was used for anesthesia. Mice were euthanized with carbon dioxide. The abscess infection model was performed as previously described (51). *K. pneumoniae* KPLN649 and *P. aeruginosa* LESB58 were grown to an OD_600_ of 1.0 in dYT broth. Prior to injection, bacterial cells were washed twice with sterile PBS and resuspended to 5 × 10^7^ CFU for *P. aeruginosa* LESB58 and 1 × 10^9^ CFU for *K. pneumoniae* KPLN49. A 50 μl bacterial suspension was injected into the right side of the dorsum. Up to 10 mg.kg^−1^ niclosamide and 5 mg.kg^−1^ colistin, each dissolved in 2.5% DMSO, 42.5% PEG400, were tested for skin toxicity prior to efficacy testing. Treatment was applied directly into the subcutaneous space into the infected area (50 μl) at 1 h post infection. The progression of the disease/infection was monitored daily and skin abscesses were excised (including all accumulated pus) on day three, homogenized in sterile PBS using a Mini-Beadbeater-96 (Biospec products, Bartlesville, OK) for 5 min and bacterial counts determined by serial dilution. Experiments were performed at least 3 times independently with 3 to 4 animals per group.

## The Appendix is comprised of Tables S1-S3 and Figures S1-S6

**Table S1.**
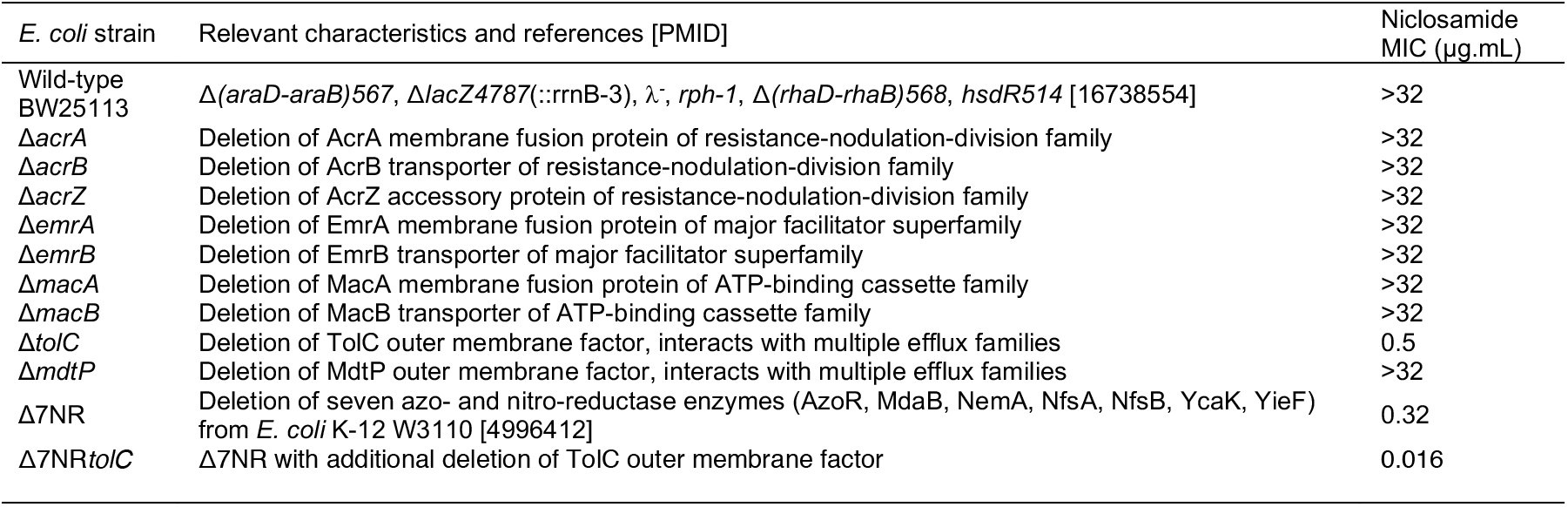
*E. coli* strains utilized in this study.

**Table S2.**
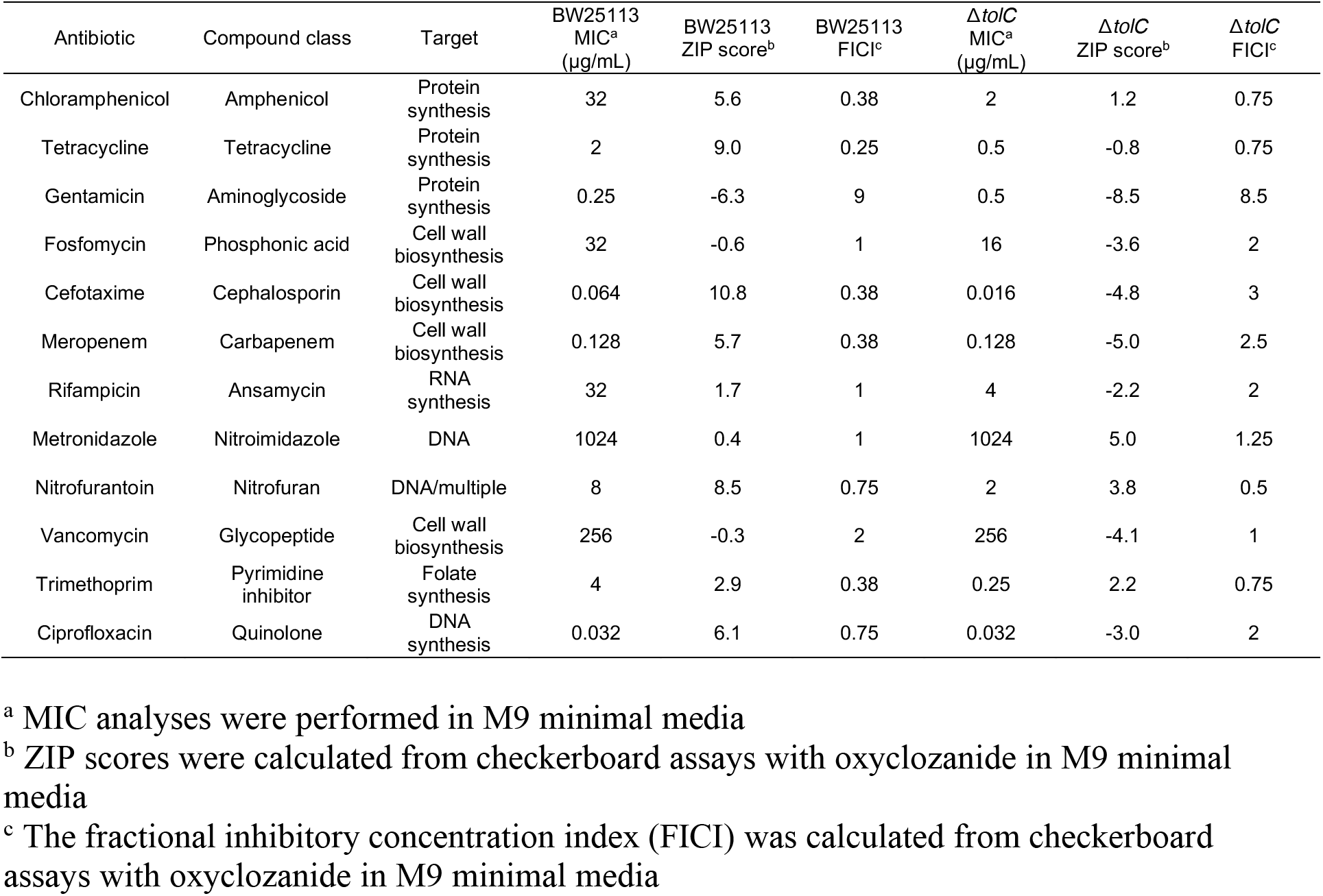
Antibiotic compounds utilized in this study.

**Table S3.** Clinical strains utilized in this study.

| Strain | Colistin MIC<br>(µg/mL) | Relevant characteristics | Reference [PMID] or source |
| --- | --- | --- | --- |
| ***Pseudomonas aeruginosa*** |  |  |  |
| *P. aeruginosa* LESB58 | 4 | Liverpool Epidemic Strain isolate | [26814180] |
| *P. aeruginosa* NZRM4034 | 1 | Ceftazidime/piperacillin resistant isolate | ESR resources |
| ***Acinetobacter baumannii*** |  |  |  |
| *A. baumannii* Ab5075 | 0.25 | Virulent wound isolate | [26991296] |
| *A. baumannii* NZRM3289 | 1 | Urine isolate (ATCC19606) | ESR resources |
| ***Klebsiella pneumoniae*** |  |  |  |
| *K. pneumoniae* KPLN49 | 0.5 | Wild-type strain | [23665232] |
| *K. pneumoniae* NZRM4387 | 1 | Extended spectrum β-lactamase urine isolate | ESR resources |
| ***Enterobacter cloacae*** |  |  |  |
| *E. cloacae* 218R1 | 0.25 | Class C β-lactamase overproducing strain | [24865555] |
| ***Escherichia coli*** |  |  |  |
| *E. coli* E38 (serotype O78:H<sup>-</sup>) | 0.25 | Human peritoneum isolate | BEI Resources |
| *E. coli* NZRM4403 | 0.5 | β-lactamase overproducing human blood isolate | ESR resources |

**Fig S1.**
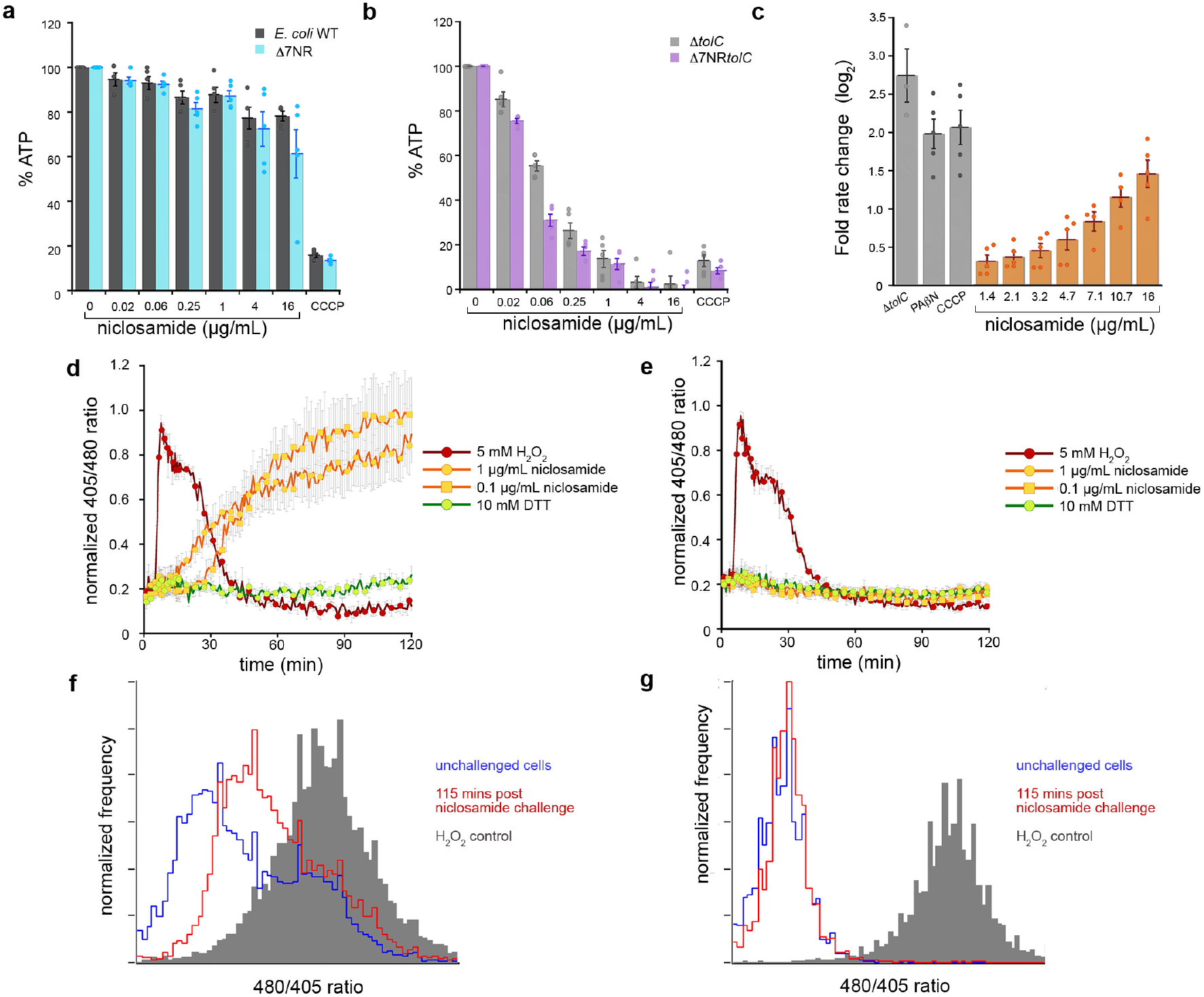
Niclosamide mechanisms of action. **a-b**, ATP concentration was investigated using a luminescent probe, BacTiter-Glo™. Following incubation for 60 min with 0.016 - 16 μg.mL^−1^ niclosamide or 32 μg.mL^−1^ CCCP, percent ATP was determined via comparison to a DMSO control in (**a**) *E. coli* BW25113 (WT) and Δ7NR strains, and (**b**) Δ*tolC* and Δ7NR*tolC* strains. **c**, Fold rate change of Hoechst H33342 fluorescence was measured over 10 min (at 355 nm and 460 nm for excitation and emission, respectively) and compared to a DMSO-only control. Δ*tolC* was employed as a disrupted efflux control. *E. coli* cells were grown in MHB media supplemented with 5 mM EDTA for permeabilization, and were administered 28 μg.mL^−1^ PAβN, 5 μg.mL^−1^ CCCP, or 1.4 to 16 μg.mL^−1^ niclosamide. Error bars indicate SEM. **d-e**, Intracellular oxidation levels were measured over 120 min in (**d**) Δ7NR*tolC* and (**e**) Δ7NR strains constitutively expressing redox-sensitive GFP (roGFP) following administration of 5 mM H_2_O_2_ (oxidized control), 1 mM DTT (reduced control), 1 μg.mL^−1^ niclosamide or 0.1 μg.mL^−1^ niclosamide. **f-g**, Histograms of the 405/480-nm ratios of intracellular redox potential of (**f**) Δ7NR*tolC* and (**g**) Δ7NRcells prior to administration of niclosamide (blue) and 115 min after administration of 1 μg.mL^−1^ niclosamide. The gray histogram represents the oxidized control (10 mM H_2_O_2_).

**Fig S2.**
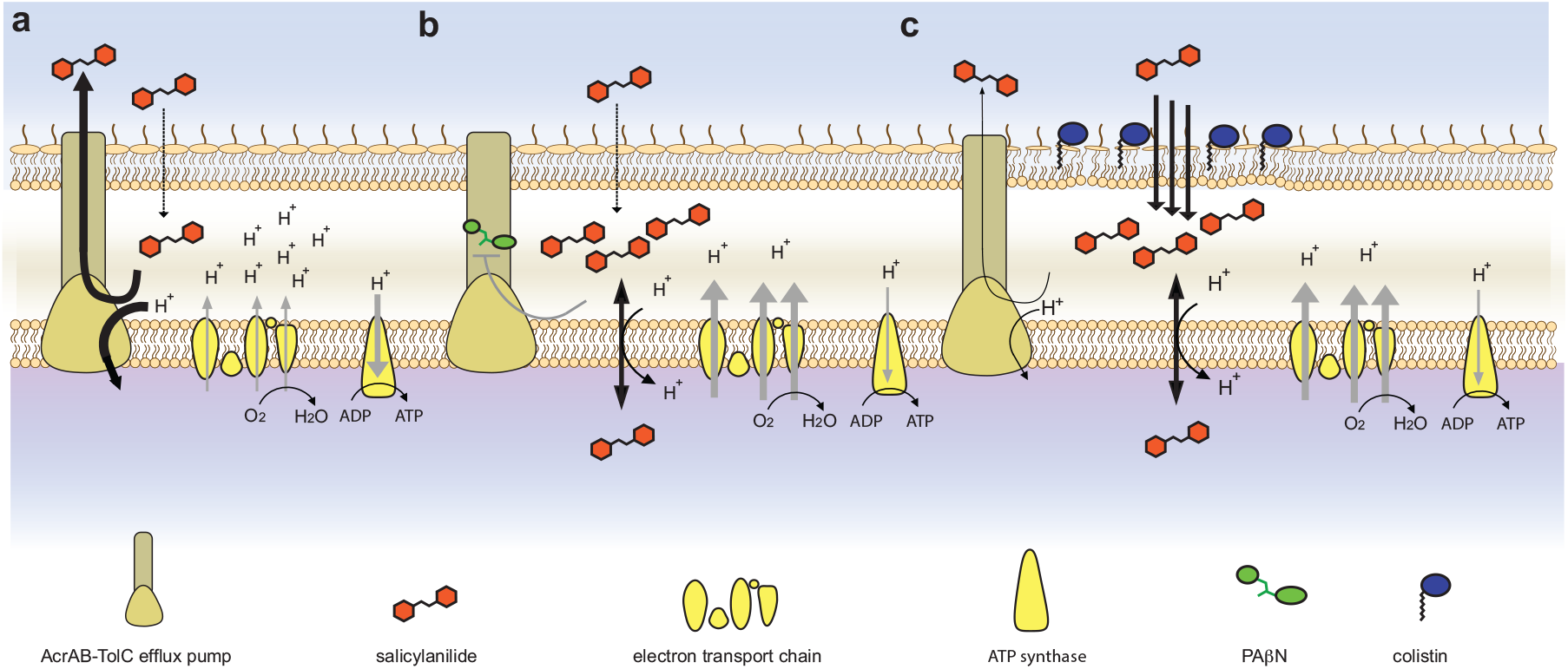
Proposed salicylanilide mechanisms of action. **a**, A salicylanilide crosses the outer membrane but is expelled from the cell *via* TolC-mediated efflux (PMF-dependent); electron transport, membrane polarization, oxygen consumption and ATP synthesis are not affected. **b**, When TolC is inhibited by compounds such as PAβN, salicylanilides uncouple the electron transport chain, dissipate the PMF, increase oxygen consumption and decrease ATP production. **c**, When the outer membrane is disrupted via compounds such as colistin, salicylanilides rapidly enter the cell, overwhelming TolC-mediated efflux, uncoupling the electron transport chain, dissipating the PMF (inhibiting PMF-dependent efflux), increasing oxygen consumption and decreasing ATP production.

**Fig. S3.**
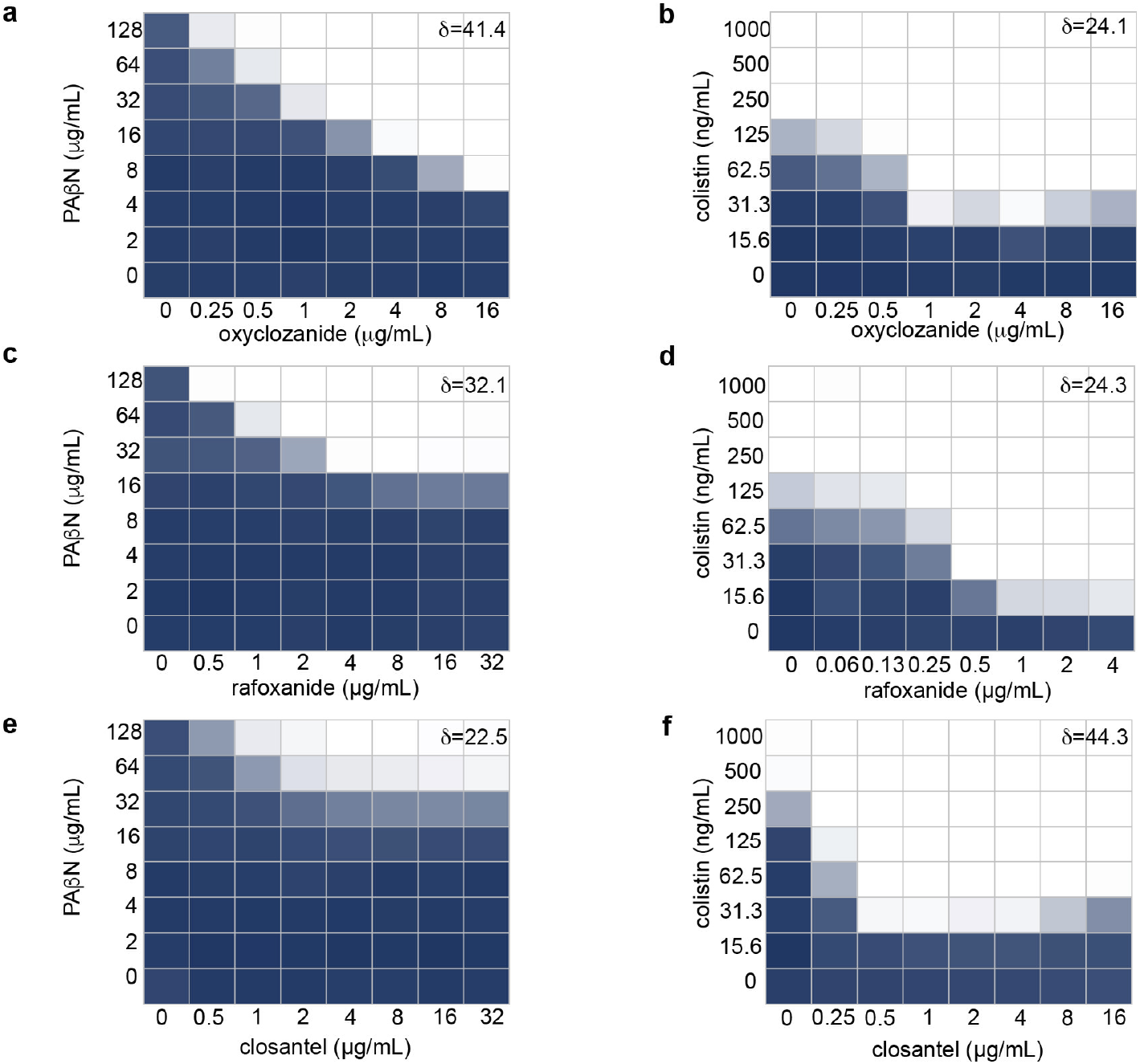
The synergistic relationships of salicylanide derivatives with PAβN and colistin. Using *E. coli* checkerboard analyses, the combined inhibitory effects of 0 to 256 μg.mL^−1^ oxyclozanide and either (**a**) 0 - 128 μg.mL^−1^ PAβN, or (**b**) 0 - 250 ng.mL^−1^ colistin; 0 - 32 μg.mL^−1^ rafoxanide and either (**c**) 0 - 128 μg.mL^−1^ PAβN, or (**d**) 0 - 250 ng.mL^−1^ colistin; and 0 - 32 μg.mL^−1^ closantel and either (**e**) 0 - 128 μg.mL^−1^ PAβN, or (**f**) 0 - 250 ng.mL^−1^ colistin were tested. Bacterial growth is shown as a heat plot. The ZIP synergy score (δ) is presented as the average interaction from the dose-response landscape. Data presented were averaged from at least 3 (typically >6) independent experiments with SEM <15%.

**Fig. S4.**
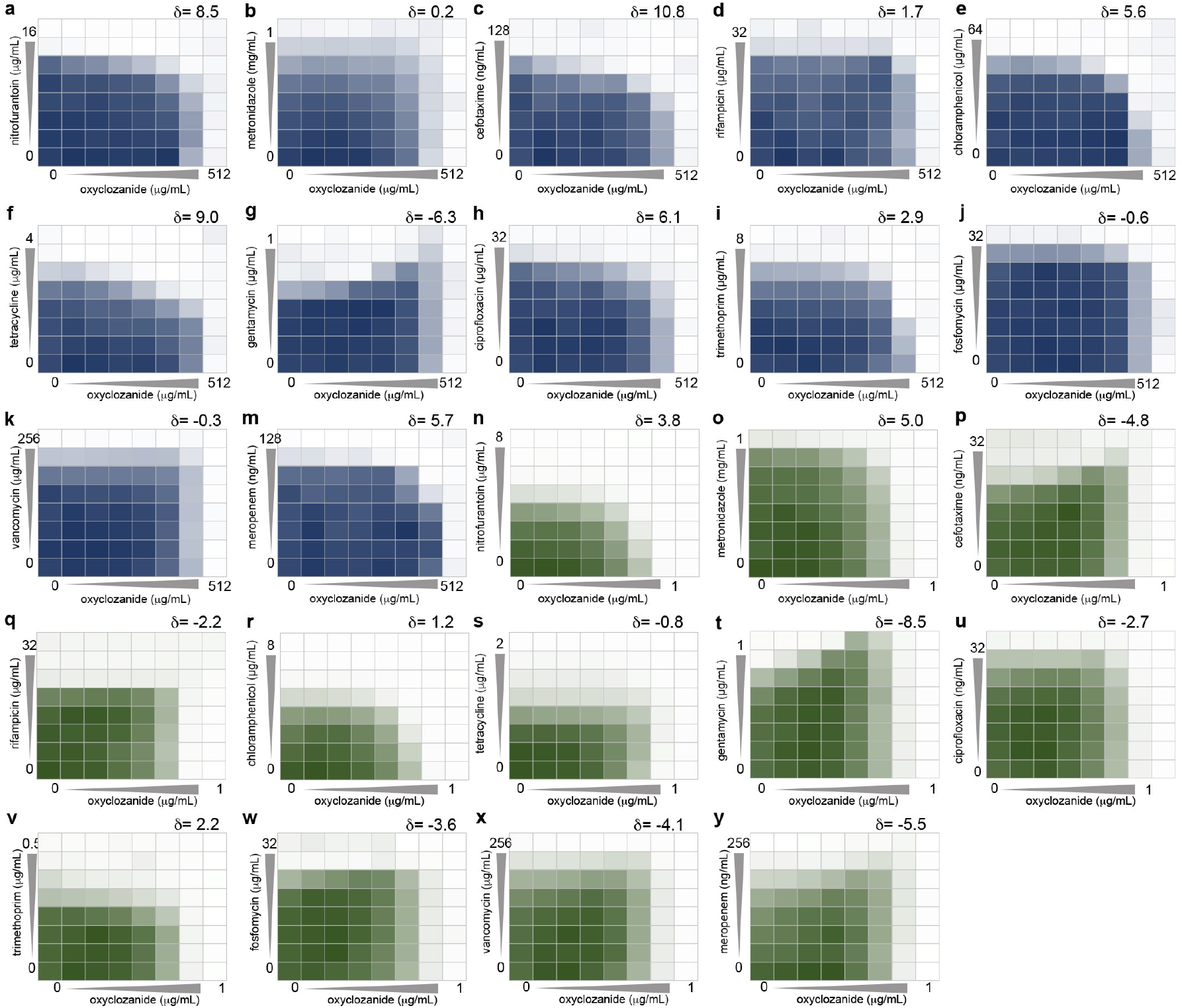
The synergistic and antagonistic relationships of oxyclozanide with diverse clinical antibiotics. **a-m**, The combined inhibitory effects of 0 to 512 μg.mL^−1^ oxyclozanide and either (**a**) 0 to 16 μg.mL^−1^ nitrofurantoin, or (**b**) 0 to 1 mg.mL^−1^ metronidazole, or (**c**) 0 to 128 ng.mL^−1^ cefotaxime, or (**d**) 0 to 32 μg.mL^−1^ rifampicin, or (**e**) 0 to 64 μg.mL^−1^ chloramphenicol, or (**f**) 0 to 4 μg.mL^−1^ tetracycline, or (**g**) 0 to 1 μg.mL^−1^ gentamicin, or (**h**) 0 to 32 ng.mL^−1^ ciprofloxacin, or (**i**) 0 to 8 μg.mL^−1^ trimethoprim, or (**j**) 0 to 32 μg.mL^−1^ fosfomycin, or (**k**) 0 to 256 μg.mL^−1^ vancomycin, or (**m**) 0 to 128 ng.mL^−1^ meropenem were tested against *E. coli* using checkerboard analyses in minimal media. Bacterial growth is shown as a heat plot. **n-y**, The combined inhibitory effects of 0 to 1 μg.mL^−1^ oxyclozanide and either (**n**) 0 to 8 μg.mL^−1^ nitrofurantoin, or (**o**) 0 to 1 mg.mL^−1^ metronidazole, or (**p**) 0 to 32 ng.mL^−1^ cefotaxime, or (**q**) 0 to 32 μg.mL^−1^ rifampicin, or (**r**) 0 to 8 μg.mL^−1^ chloramphenicol, or (**s**) 0 to 2 μg.mL^−1^ tetracycline, or (**t**) 0 to 1 μg.mL^−1^ gentamicin, or (**u**) 0 to 32 ng.mL^−1^ ciprofloxacin, or (**v**) 0 to 0.5 μg.mL^−1^ trimethoprim, or (**w**) 0 to 32 μg.mL^−1^ fosfomycin, or (**x**) 0 to 256 μg.mL^−1^ vancomycin, or (**y**) 0 to 256 ng.mL^−1^ meropenem were tested against *E. coli* Δ*tolC* using checkerboard analysis in minimal media. Bacterial growth is shown as a heat plot. The ZIP synergy score (δ) is presented as the average interaction from the dose-response landscape. Data presented were averaged from at least 3 (typically >6) independent experiments with SEM <15%.

**Fig. S5.**
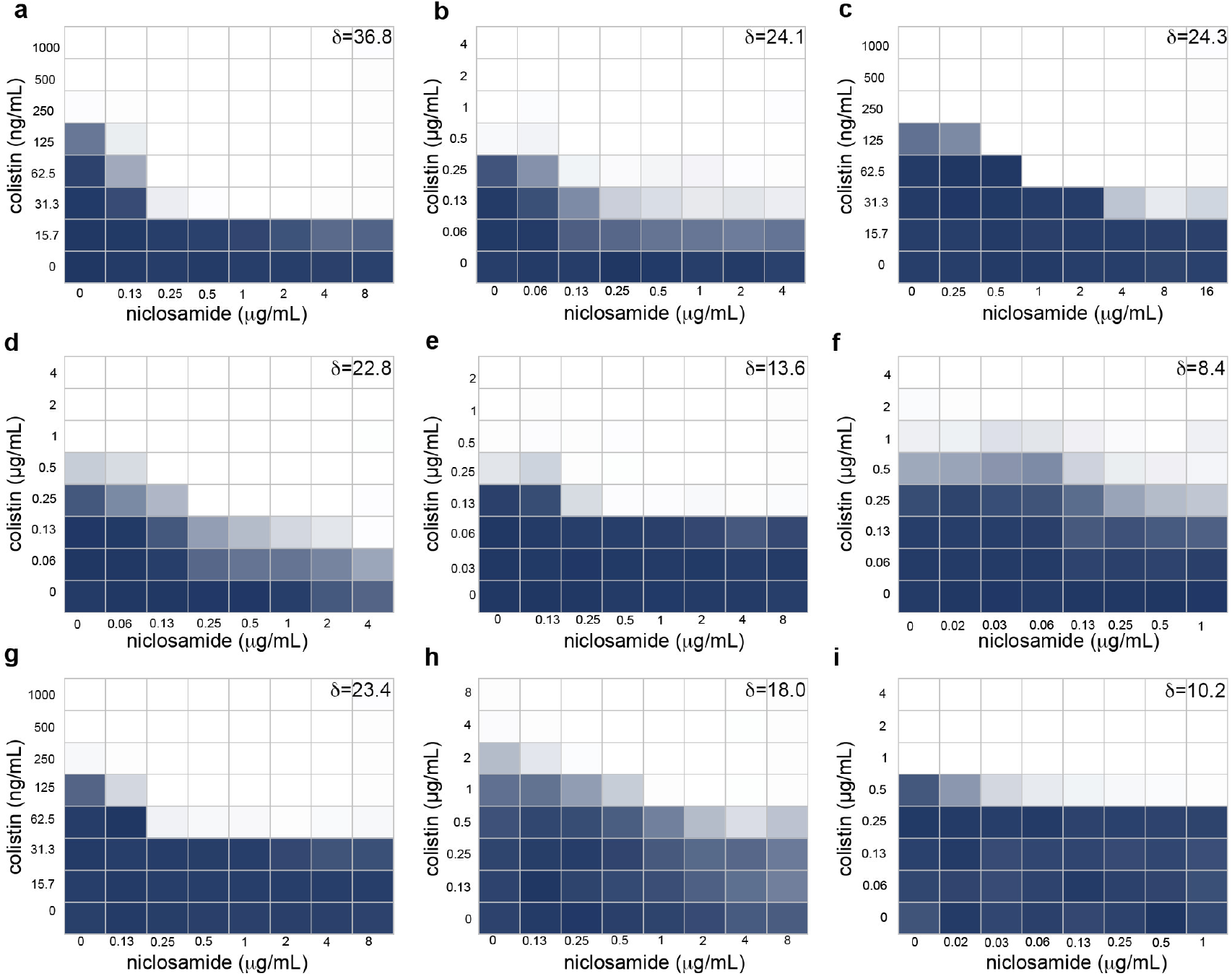
The synergistic relationships of niclosamide and colistin against clinical isolates. **a-i**, The combined inhibitory effects of up to 16 μg.mL^−1^ niclosamide and up to 8 μg.mL^−1^ colistin were tested using checkerboard analysis against (**a**) *E. coli* E38, or (**b**) *E. coli* NZRM4403, or (**c**) *A. baumannii* Ab5075, or (**d**) *A. baumannii* NZRM3289, or (**e**) *K. pneumoniae* KPLN649, or (**f**) *K. pneumoniae* NZRM4387 or (**g**), *E. cloacae* 218R, or (**h**) *P. aeruginosa* LESB15, or (**i**) *P. aeruginosa* NZRM4034. The ZIP synergy score (δ) is presented as the average interaction from the dose-response landscape. Data presented were averaged from at least 3 independent experiments with SEM <18%.

**Fig. S6.**
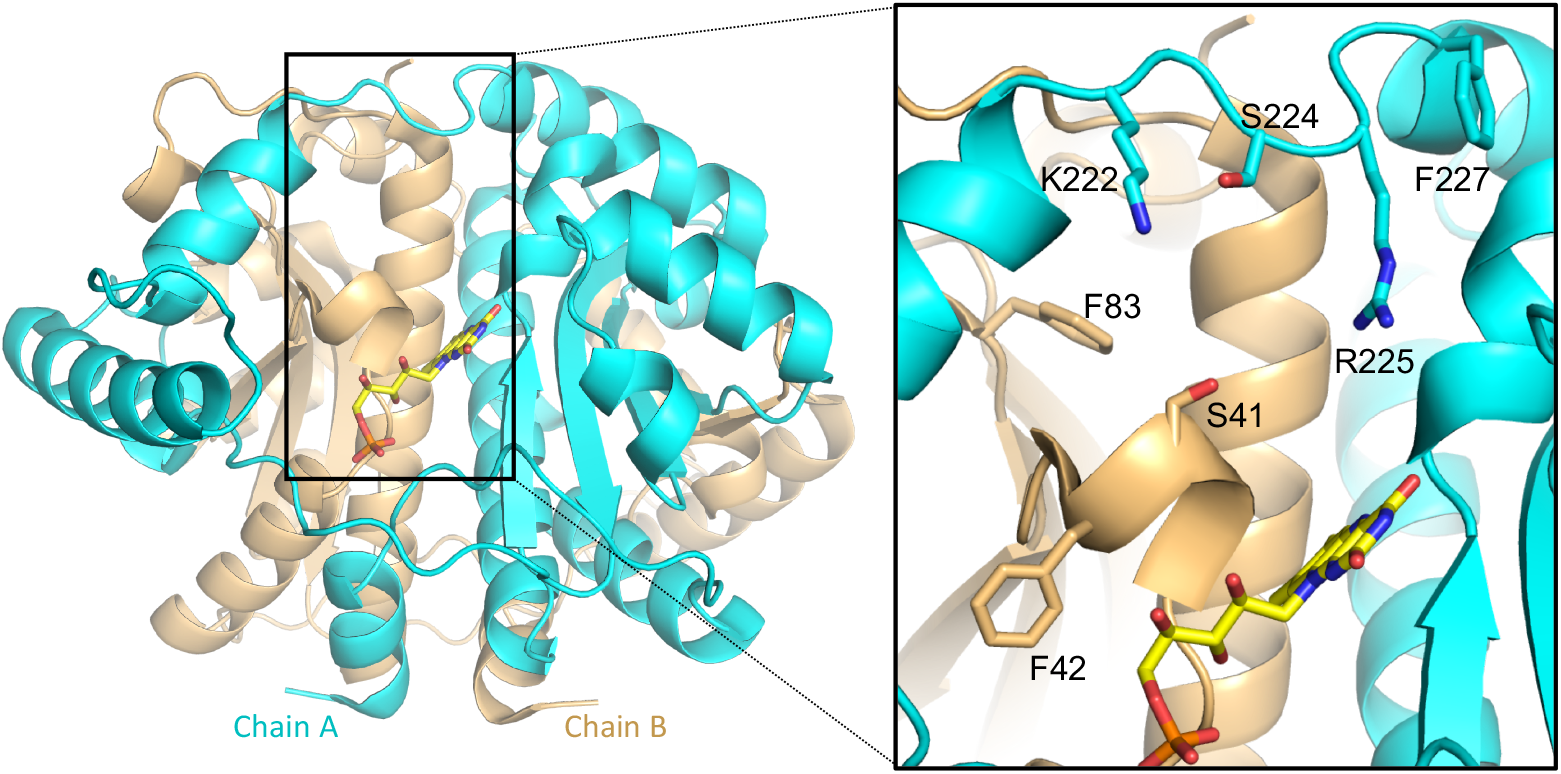
Combinatorial mutagenesis of *E. coli* NfsA. A ribbon diagram displays the dimeric structure of PBD ID: 1f5v (*E. coli* NfsA). Monomers are colored in cyan or gold respectively. FMN is depicted as a stick model with carbons colored in yellow; one active site is shown for clarity. ***Inset:*** The NfsA active site. Residues that were randomized to generate the NfsA variant library are labeled and displayed in stick models.

